# Priming of MSCs with inflammation-relevant signals affects extracellular vesicle biogenesis, surface markers, and modulation of T cell subsets

**DOI:** 10.1101/2020.04.30.066456

**Authors:** Seth Andrews, Ty Maughon, Ross Marklein, Steven Stice

## Abstract

Although considerable evidence exists supporting the use of mesenchymal stromal cells (MSCs) for treating immune diseases, successful clinical translation has been challenging and has led researchers to investigate cell-free alternatives. MSC-derived extracellular vesicles (MSC-EVs) have been shown to mediate a significant portion of the observed therapeutic effect, including immunosuppression. MSCs have been shown to respond to different aspects of the injury microenvironment such as inflammatory cytokines and hypoxia, although acidosis has not been investigated and different conditions have not been assessed in terms of their effects on MSC-EV function. This study investigated the effects of acidosis, hypoxia, and inflammatory cytokine priming on MSCs and MSC-EVs. We cultured MSCs in the presence of acidosis, hypoxia, or inflammatory cytokines (Interferon-gamma and Tumor Necrosis Factor-alpha) and compared the characteristics of their EVs as well as their uptake by and suppression of different T cell subsets. MSCs showed a greater effect on suppressing activated CD4^+^ and CD8^+^ T cells than MSC-EVs. However, MSC-EVs from MSCs primed with acidosis increased CD4^+^ and CD8^+^ regulatory T cell frequency in vitro. This functional response was reflected by MSC-EV uptake. MSC-EVs from acidosis-primed MSCs were taken up by CD4^+^ and CD8^+^ regulatory T cells at a significantly higher level than MSC-EVs from control, hypoxic, and inflammatory cytokine groups. These data suggest that a simple low-cost alteration in MSC culture conditions, acidosis, can generate extracelluar vesicles that have a desirable influence on anti inflammatory T cell subtypes.

## 1. Introduction

Mesenchymal stromal cells (MSCs) are being explored as an immunomodulatory therapy to treat immune diseases such as osteoarthritis, multiple sclerosis, and Parkinson’s disease (1, 2); however, cell therapies face challenges associated with manufacturing consistent, high quality products. Although promising as an allogeneic ‘off-the-shelf’ therapy, cryopreserved and thawed MSCs have diminished efficacy and require a recovery period at point of care when compared to fresh, non-thawed MSCs (3). Transplantation of any cell therapy, even MSCs, can also raise safety concerns in the event of uncontrolled differentiation into undesired tissue or promotion of tumor growth following engraftment (4–6).

Cell-free extracellular vesicle (EV) preparations from MSC cultures that possess similar MSC therapeutic function are a potential alternative to MSC therapy. The MSC secretome is responsible for much of their regenerative and immunomodulatory functions (7). MSC immunomodulation via secreted factors has been directly linked to T cell suppression, as shown in a transwell system that prevents cell-cell contact between MSCs and T cells (8–10). Extracellular vesicles are nanoscale vesicles released from all cell types and participate in intercellular signaling through transference of bioactive molecules including RNA, proteins, and lipids (11). MSC-EVs do not divide, engraft, or dynamically respond to their environment like MSCs, thus addressing concerns with tumorigenicity and ectopic tissue growth (5, 6). In vivo data from our lab indicates that some EVs can suppress systemic levels of inflammatory T cells after injury in mice (12). MSC-EVs have demonstrated preliminary evidence of regenerative effects ranging from recovery from myocardial ischemia and reperfusion injury (13), stroke (14), gentamicin induced acute kidney injury (15), and allogeneic skin grafts (16).

The injury microenvironment is often characterized by inflammation, involving local hypoxia, acidosis, as well as the presence of cytokines such as Tumor Necrosis Factor alpha (TNF-α) and Interferon gamma (IFN-γ) (17–19). MSCs are known to respond to priming by inflammatory environments by switching to an “activated” immunosuppressive phenotype. This often involves upregulating expression of regenerative and antiinflammatory factors such as Vascular Endothelial Growth Factor, Indolamine 2,3-Dioxygenase, Transforming Growth Factor Beta, and Prostaglandin E2 (20). More recently, inflammatory and hypoxic priming have been shown to increase the potency of MSC-EV immunomodulation (21–23). EVs derived from MSCs primed with hypoxia were more effective than EVs from non-primed MSCs in inducing macrophage proliferation and type 2 macrophage polarization (21). TGF-β and IFN-γ primed MSCs produced EVs that were more effective in inducing regulatory T cell (T_reg_) formation than those from resting MSCs (23). This suggests that MSC-EVs are involved in the MSC response to inflammatory priming. However, acidic priming has not been investigated in this manner, and different cell culture conditions have not been compared within the same study.

To determine the effects of cell culture preconditioning (i.e. priming) on MSC-EV quality and their potential role in a therapeutically-relevant function (T cell suppression) we performed the following studies. First, we examined the effects of hypoxia, acidosis, and inflammatory cytokines on MSC-EV biogenesis and release as compared to unprimed MSC culture conditions. Activation of different T cell subsets treated with both MSCs and MSC-EVs was comprehensively profiled. Inflammatory cytokine priming increased EV size, and decreased relative expression of a panel of surface markers. Meanwhile, acidosis and hypoxia increased EV yield while having little effect on their surface marker composition. Additionally, while MSCs in direct contact demonstrated greater suppression of effector T cells than MSC-EVs, MSC-EVs derived from MSCs primed with acidosis induced the formation of T_regs_ while other MSC-EV groups had no significant effect on T cell activation. Therefore, precise control and monitoring of the pH during MSC-EV manufacturing should be further explored as a means to effect MSC-EV immunomodulatory function.

## 2. Methods

### 2.1 Cell culture and priming

Human female wharton’s jelly MSCs (Lifeline Cell Technologies) referred to as MSC here, were plated at 5000 cells/cm^2^ on tissue culture flasks in complete medium (Alpha-Minimum Essential Medium (Gibco), 10% defined fetal bovine serum (Hyclone), 2 mM L-glutamine, 50 U/mL penicillin, 50 μg/mL streptomycin (Gibco) and allowed to grow to 80% confluence (20,000–25,000 cells/cm^2^). They were harvested using 0.05% trypsin (Gibco) and replated at 5000 cells/cm^2^. All proliferation cultures were maintained at 37°C and 5% CO_2_. All cells used in experiments had undergone fewer than 10 passages and had over 90% viability at harvest as assessed by Trypan blue staining.

Several environments were used to prime MSCs when they reached 80% confluence (Fig. S1). Metabolic acidosis was induced through the addition of HCl to complete media to lower the pH to 7.1 ± 0.05 (24). These MSCs were termed LPH-MSCs, with their EVs being LPH-EV. A hypoxic environment was created by placing the cell culture vessels in a hypoxia incubator chamber (STEMCELL Technologies, Cambridge MA), which was then filled with a gas mixture containing 2% O_2_, 5% CO_2_, and 93% N_2_ (Airgas, Radnor PA) for 5 minutes at 2 psi according to the manufacturer’s recommendation (25, 26). The MSCs undergoing this priming were termed LO2-MSCs, and their EVs were LO2-EV. An inflammatory environment was created by adding the cytokines TNF-α and IFN-γ (Sigma, Burlington MA) to the medium at 15 and 20 ng/mL, respectively. These MSCs were termed INF-MSCs, with their EVs being INF-EV. These environments were used separately to precondition MSCs for 48 hours at 37°C prior to receiving EV isolation media or being placed in co-culture with human Peripheral Blood Mononuclear Cells (PBMCs). A final group of MSCs remained in complete growth medium at pH 7.4 and 20% oxygen as described above for 48 hours. The normal culture MSCs and their EVs were NC-MSC and NC-EV, respectively.

PBMCs (STEMCELL Technologies, Cambridge MA) were thawed into RPMI media (RPMI 1640, 10% FBS, 50 U/mL penicillin, 50 μg/mL streptomycin) and cultured for 16hrs at 37°C and at 5% CO_2_ prior to use in experiments.

### 2.2 EV isolation and characterization

After priming, MSCs were rinsed twice with PBS before adding fresh serum free medium (Alpha-Minimum Essential Medium (Gibco), 2 mM L-glutamine, 50 U/mL penicillin, 50 μg/mL streptomycin (all from Gibco/Invitrogen)) and incubating cultures for 24 hours. The resulting conditioned media were collected and passed through 0.22 μm filters to remove cells and large debris. The media were subjected to ultrafiltration with a 100kDa MWCO (Amicon, Millipore-Sigma, Burlington MA) at 4000g for 10 minutes as we previously published (27). The EVs were part of the retentate and were then washed twice with PBS +/+ (Thermo Fisher Scientific, Waltham, MA) at 2000g for 10 minutes. The EVs in PBS+/+ were then collected, aliquoted, and frozen at −20°C.

For each EV isolation, nanoparticle tracking analysis (NTA) was performed using a Nanosight NS3200 (Nanosight, Salisbury UK) according to the manufacturer’s recommendations. Briefly, aliquots of EV suspensions were thawed at room temperature and diluted to 10^7^-10^9^ particles/mL with the same lot of PBS +/+ the EVs were isolated in. A minimum of three samples and five one-minute videos were recorded for each EV isolation. All videos were captured at the same camera level and analyzed with the same detection threshold.

The size and size distribution of vesicles was further verified via Dynamic Light Scattering using a Malvern Zetasizer Nano ZS Analyzer (Malvern Instruments, Malvern, UK). Samples were diluted to a total vesicle concentration of approximately 2×10^8^ vesicles/ml in 0.22μm filtered Phosphate Buffered Saline containing Calcium and Magnesium, pH 7.4 prior to measurements. Disposable polystyrene cuvettes were rinsed with 1mL of filtered PBS +/+ prior to adding sample. Measurements were taken using cuvette with 800uL of prepared sample.

**Supplementary Figure 1:**
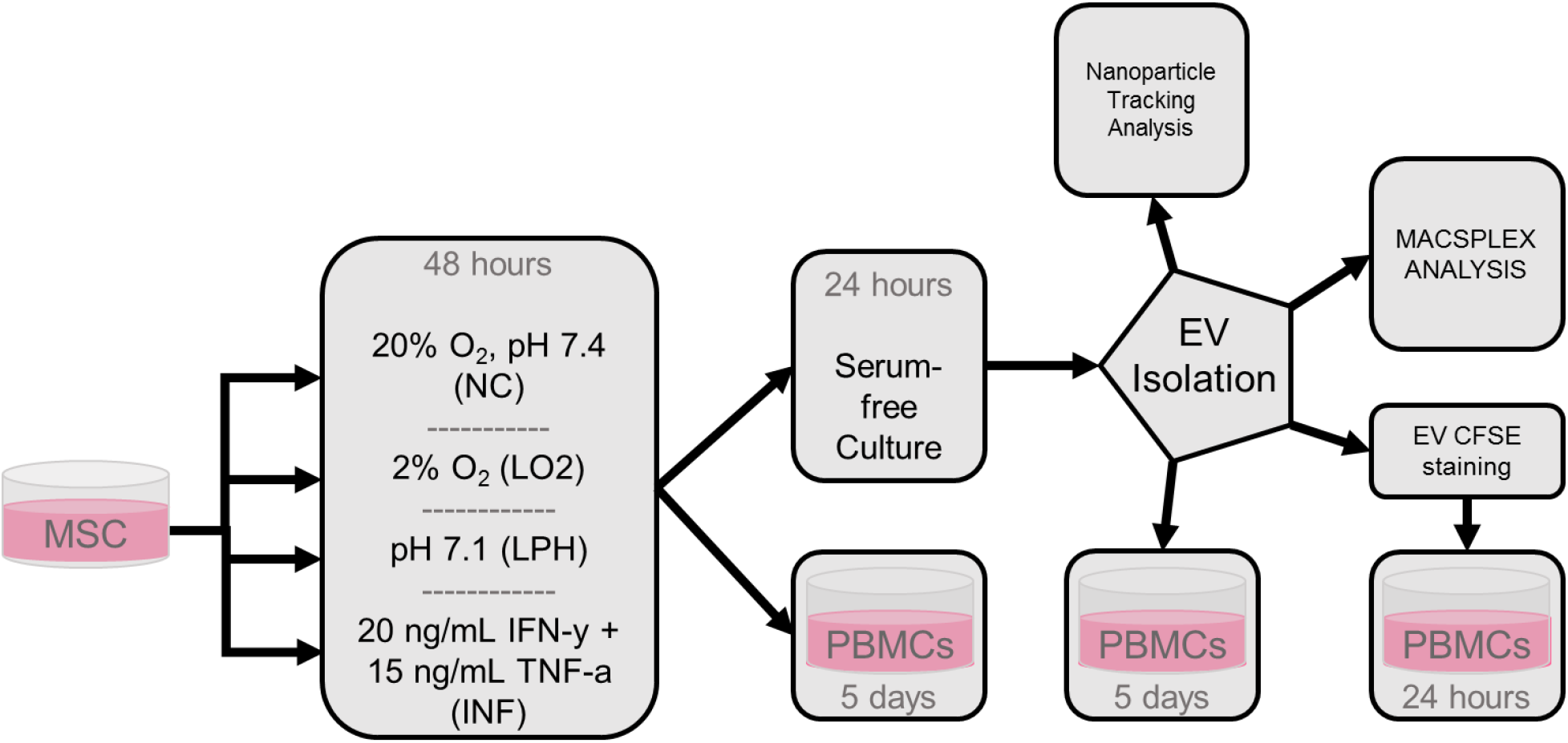
Experimental workflow. MSCs were split into groups that each underwent different priming steps. MSCs were then either co-cultured with PBMCs for 5 days or used for EV isolation. Isolated EVs were used in NTA or MACSPLEX analysis, stained with CFSE and incubated for 24 hours with PBMCs, or incubated for 5 days with PBMCs.

EV surface marker characterization was performed using the MACSPLEX Exosome Kit (Miltenyi Biotec, Bergisch Gladbach, Germany) according to the manufacturer’s directions. Briefly, an equal number of EVs as determined by NTA were analyzed from each isolation in triplicate. Flow cytometry analysis was performed using a CytoFLEX S (Beckman Coulter, Hialeah, Florida) alongside bead only controls, with FlowJo (Ashland, OR) being used for data analysis. Data was processed with background subtraction and normalized to the median of the average value of CD9, CD63, and CD81 for each sample (28). The data was transformed to be a percentage of the difference between the maximum and minimum relative expression of a marker. Principal component analysis (PCA) was performed on the transformed data using JMP (SAS Institute, Cary NC).

### 2.3 EV uptake assay

PBMCs were added to each well at 500,000 cells per well in 48 well plates. Stimulating anti-CD3/CD28 Dynabeads (Thermo Fisher Scientific, Waltham, MA) were then added at 500,000 per well. EVs were stained with CFSE (Thermo Fisher Scientific, Waltham, MA) following a protocol modified from Morales-Kastresana (29). 40 μM CFSE in PBS +/+ was added to an equivalent volume of EVs and incubated for 2 hours in the dark at 37°C. Excess dye was quenched with an equivalent volume of 0.1 % BSA, and the whole mixture was rinsed with PBS +/+ and concentrated via ultrafiltration as in the initial EV isolation. 10^9^ CFSE-EVs were then added to the appropriate wells.

The assays took place in complete RPMI medium formulated as above, but with EV-depleted FBS. EVs were depleted by centrifuging FBS at 100,000g for 1 hour at 4°C (Sorvall WX Ultra 80, Thermo Fisher Scientific, Waltham, MA) and using the supernatant (30). The cultures incubated for 24 hours at 37°C, 5% CO_2_ for the duration of the experiments.

Following incubation, the PBMCs were harvested and stained for flow cytometry using Pacific Blue anti-CD4, APC anti-CD8, Brilliant Violet 711 anti-CD25, and PE anti-FOXP3. Antibodies and clones are listed in Table S2. PBMCs were first washed, then stained with Zombie Yellow viability dye, blocked with 2% FBS and Fc receptors blocked with Trustain FcX. The PBMCs were then stained for CD4, CD8, and CD25 as appropriate at room temperature in the dark for 30 minutes and fixed in FOXP3 TrueNuclear fix before storing overnight in the dark at 4°C. PBMCs were permeabilized with TrueNuclear permeabilization buffer and stained with PE anti-FOXP3 according to the manufacturer’s directions. Samples were resuspended in 2% FBS at 4°C in the dark for up to 2 days before flow analysis. All antibodies and reagents were from Biolegend (San Diego, CA) unless otherwise specified and were used at previously titrated optimal concentrations.

### 2.4 Immunomodulation assay

MSCs were plated at 20,000 cells/cm^2^ in 48 well plates in complete medium and allowed to adhere for 24 hours. The cells were then subjected to priming as previously described, followed by two PBS −/− washes.

PBMCs were labeled with CFSE (Thermo Fisher Scientific, Waltham, MA) according to the manufacturer’s instructions and 500,000 PBMCs were added to each well. After the addition of PBMCs, stimulating anti-CD3/CD28 Dynabeads (Thermo Fisher Scientific, Waltham, MA) were added at 500,000 each per well, and 10^9^ EVs were added to the appropriate wells.

The assays took place in complete RPMI medium formulated as above, but with EV-depleted FBS. EVs were depleted by centrifuging FBS at 100,000g for 1 hour at 4°C (Sorvall WX Ultra 80, Thermo Fisher Scientific, Waltham, MA) and using the supernatant (Li 2017).The cultures incubated for 5 days at 37°C, 5% CO_2_ for the duration of the experiments.

Following incubation, the PBMCs were harvested and analyzed by flow cytometry using one of two panels of conjugated antibodies. Panel 1 was composed of Pacific Blue antiCD4, APC anti-CD8, Brilliant Violet 711 anti-CD25, and PE anti-FOXP3. Panel 2 included Pacific Blue anti-CD4, APC anti-CD8, PE anti-IFN-γ, and Brilliant Violet 711 anti-TNF-α. Antibodies and clones are listed in Table S2. PBMCs were first washed, then stained with Zombie Yellow viability dye, blocked with 2% FBS and FC receptors blocked with Trustain FcX. The PBMCs were then stained for CD4, CD8, and CD25 as appropriate at room temperature in the dark for 30 minutes and fixed in 4% PFA for Panel 2 or FOXP3 TrueNuclear fix for Panel 1 before storing overnight in the dark at 4°C. Panel 1 was then permeabilized with TrueNuclear permeabilization buffer and stained for FOXP3 according to the manufacturer’s directions. Panel 2 was permeabilized with Permwash (BD Biosciences) and stained for IFN-γ and TNF-α. Samples were resuspended in 2% FBS at 4°C in the dark for up to 2 days before flow analysis. All antibodies and reagents were from Biolegend (San Diego, CA) unless otherwise specified and were used at previously titrated optimal concentrations.

**Supplementary Figure 2:**
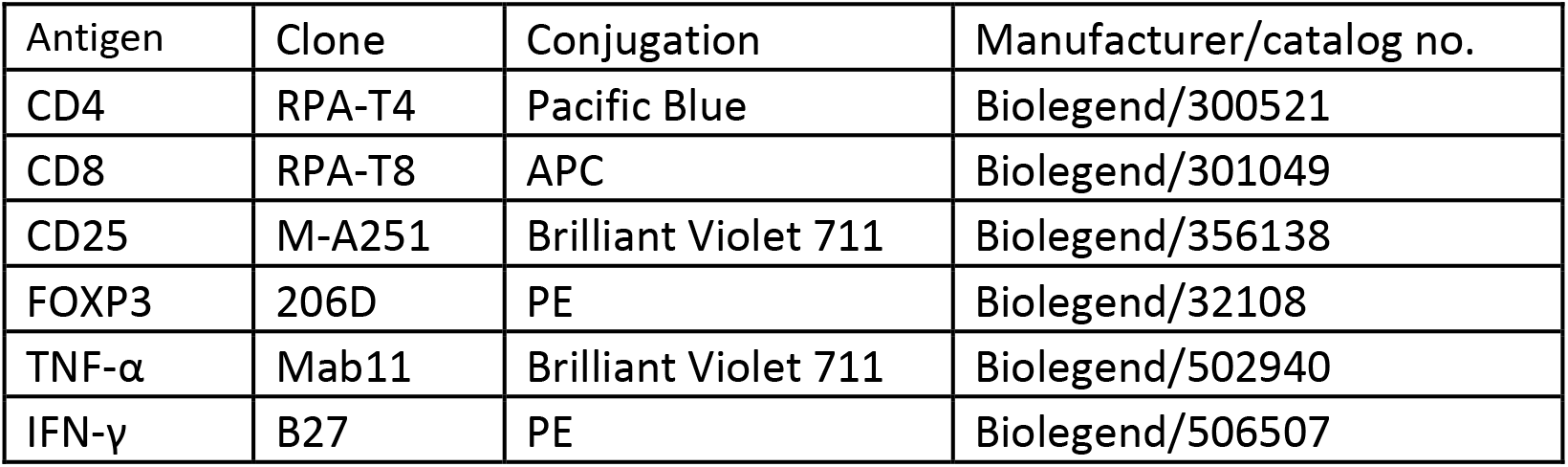
Antibody table. All antibodies used in this study, which were titrated to optimal concentration prior to experiments.

### 2.5 Flow Cytometry

All flow analysis was performed using a CytoFLEX S (Beckman Coulter, Hialeah, Florida), with 20,000 events collected per sample for the uptake and immunosuppression assays (S3). All data was analyzed using FlowJo software (Treestar, Inc., Ashland, Oregon). Cellular debris, activating beads, and doublets were gated out via scatter properties. Single-stain controls were used to generate compensation matrices, and Fluorescence-minus one controls were used to determine positive populations of CD4, CD8, CD25, TNF-α, IFN-γ, and FOXP3. Example scatter plots for the gating strategy are shown in Fig. S3.

**Supplementary Figure 3:**
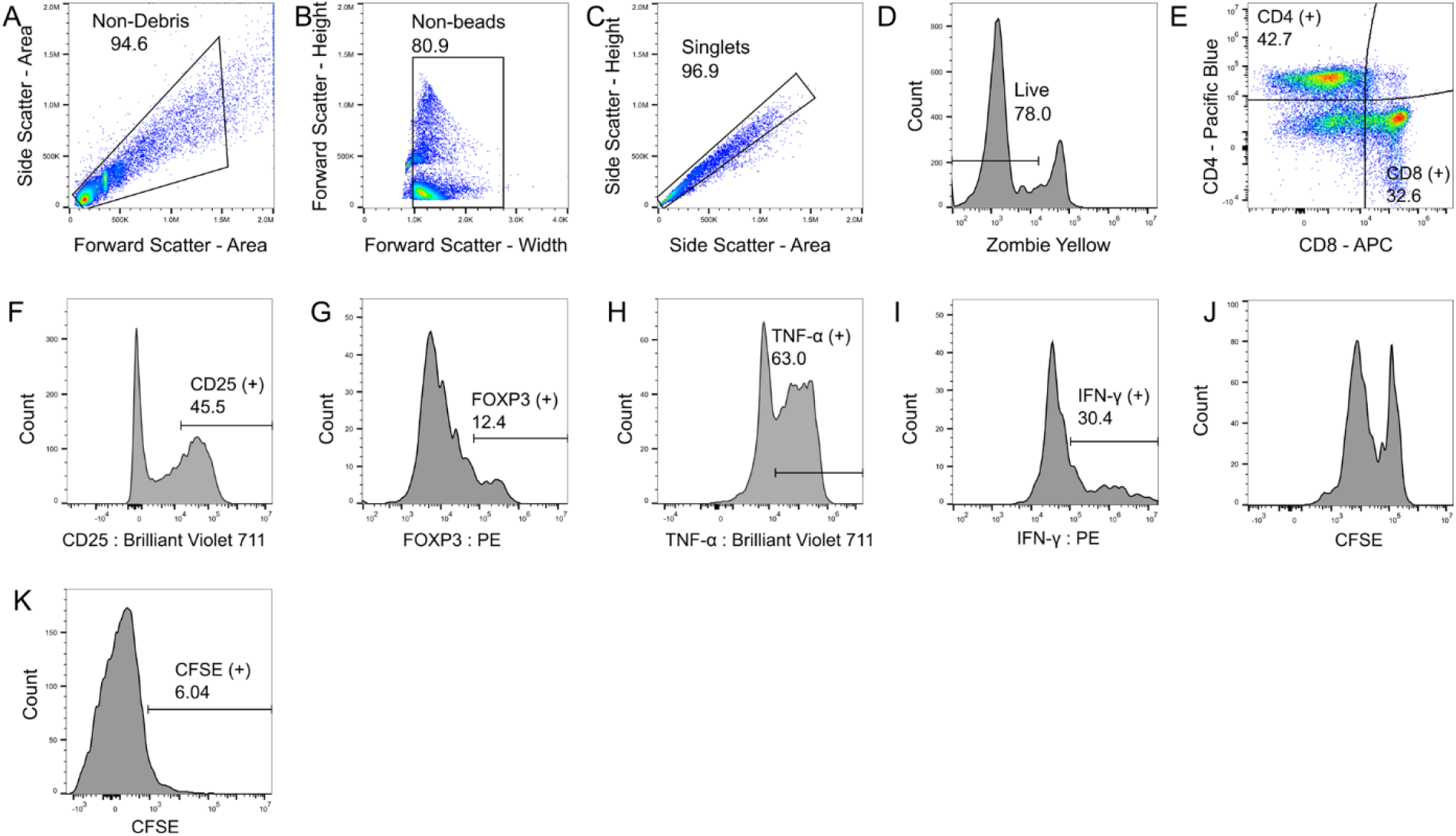
Flow analysis diagram. After successively gating out debris (A), activating beads (B) and doublets (C), live PBMCs (D) were gated based on CD4 and CD8 expression (E). Single-positive cells were then analyzed based on the panel they were stained with. Panel 1 was first gated by CD25 expression (F), then CD25^+^ cells were gated by FOXP3 expression (G). Panel 2 was gated separately by TNF-α and IFN-γ. The median fluorescence intensity of CFSE (J) was calculated for all samples to be used in proliferation calculations. For EV uptake, the populations from Panel 1 were determined at 24 hours post-treatment and T-cell subpopulations were gated by CFSE intensity (K). All gates were determined by FMO controls after compensation.

All activation parameters of T cells including CD25, FOXP3, TNF-α, and IFN-γ expression were normalized according to the formula below, so that fully activated samples (activated PBMCs with no treatment) have an average value of 100 and fully suppressed samples (resting PBMCs) have an average value of 0 (31).

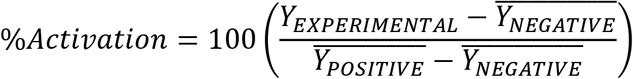

T cell proliferation was calculated according to the formula below, where MI is the median fluorescence intensity of CFSE stained samples, and PS is the proliferation score (32, 33):

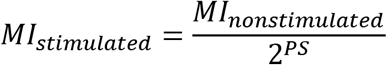

### 2.6 Statistics

All data is expressed as mean +/− SEM, with all experiments performed in triplicate unless stated otherwise. All statistical tests were one-way ANOVA against controls unless stated otherwise with Dunnett’s post-hoc test using Prism (Graphpad, San Diego CA).

## 3. Results

### 3.1 Effects of priming on MSC-EV yield and size distributions

NC-EV, LO2-EV, and LPH-EV had unimodal size distributions centered at approximately 100nm (Fig. 1A-C). The size distribution of INF-EV was bimodal, with one population centered at 100 nm as well as a larger diameter population between 150 and 400 nm (Fig. 1D). INF-EVs had a significantly higher mean diameter than NC-EVs (p<0.0001), while LPH-EV and LO2-EV did not differ in size from the control (Fig. 1E).

**Figure 1:**
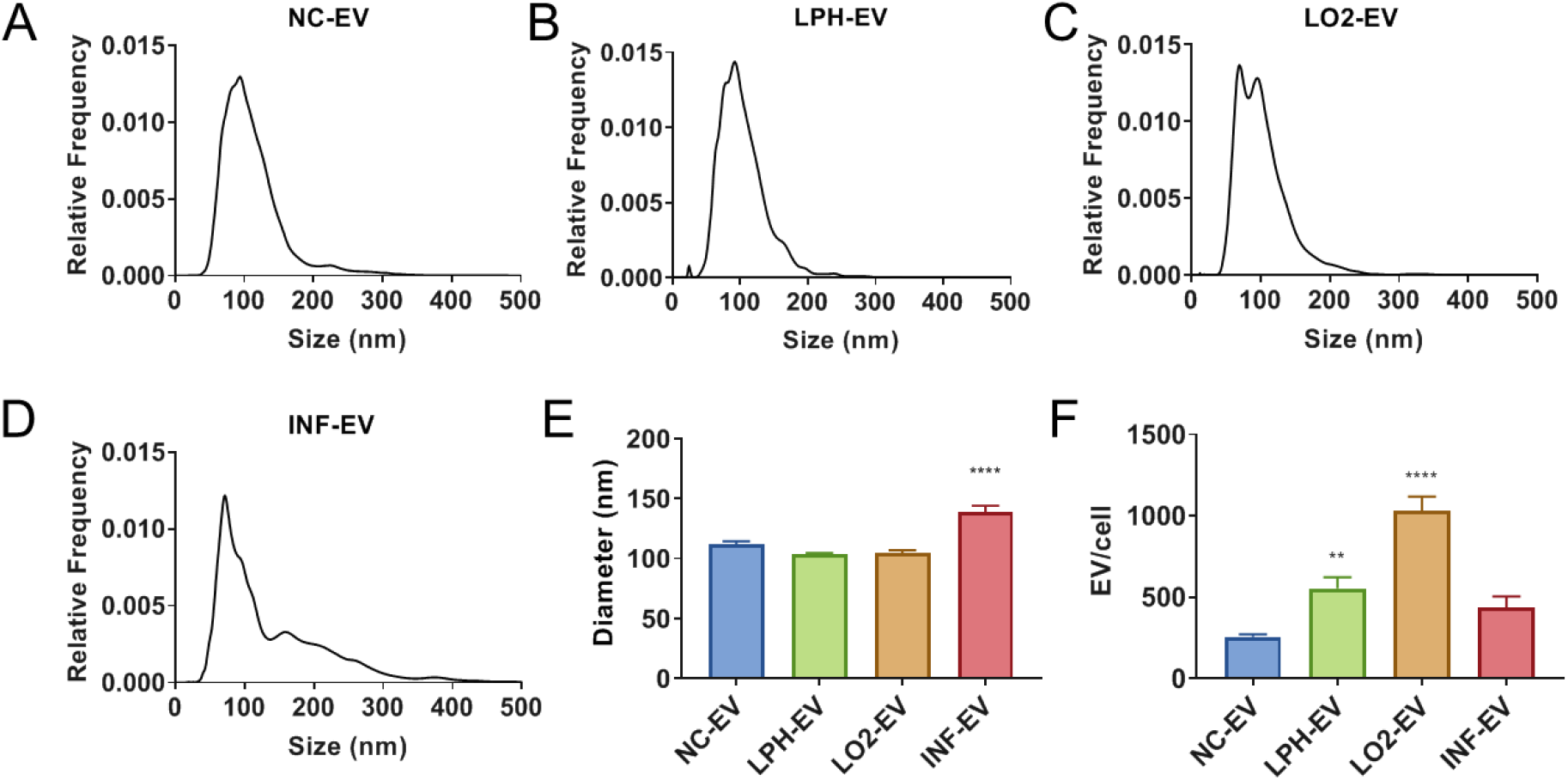
EV characterization: Size and concentration of MSC-EVs is affected by priming. Hypoxic and acidic preconditioning increased EV release. (A) Size distributions of EV groups as measured by NTA (B): EV diameter across MSC culture conditions as determined by NTA. (C): EV concentration across MSC culture conditions as determined by NTA. One-way ANOVA. Data is presented as means ± SEM. N = 3 independent experiments. (*,**, ***,****) indicate significant difference from Normal Culture at p < 0.05, 0.01, 0.001, 0.0001 by Dunnett’s post-hoc test.

Dynamic Light Scattering (DLS) analysis was used to confirm the presence of a larger population in the INF-EV group and generated similar results to NTA, despite interference from the bimodal distribution (S4). NTA analysis also revealed significantly higher EV release per cell from LO2-MSC (p<0.0001) and LPH-EV (p = 0.0028) compared to NC-MSC (Fig. 1F). Overall, hypoxic and acidic priming increased EV release, while only inflammatory priming affected EV size.

**Supplementary Figure 4:**
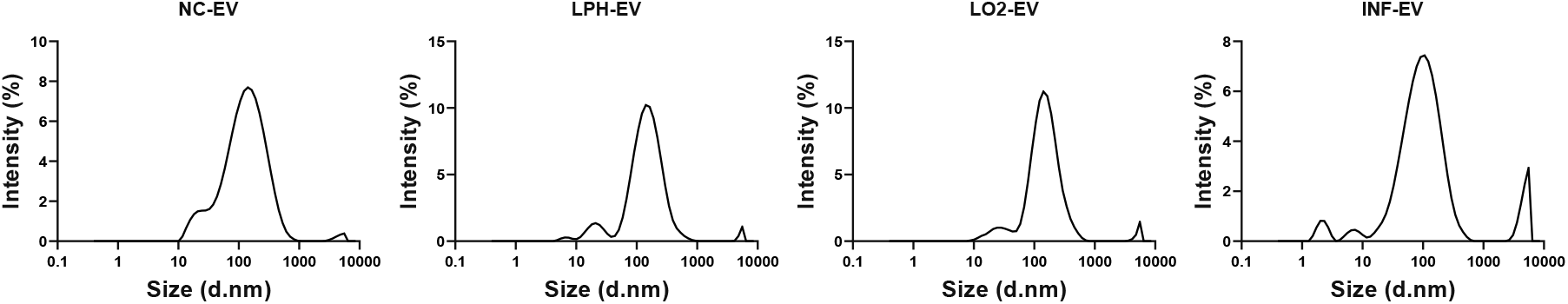
Dynamic Light Scattering of EVs. Histograms of EV size distribution confirming the existence of a large-diameter population present in INF-MSC-EV at a high concentration.

MSC (Fig. 1F). Overall, hypoxic and acidic priming increased EV release, while only inflammatory priming affected EV size.

### 3.2 Inflammatory priming affects MSC-EV surface marker expression

Surface marker characterization via MACSPLEX analysis revealed significant changes in the relative expression of EV surface markers in the preconditioned groups compared to the NC-EV group (Fig. 2A). It is important to note that the MACSPLEX assay does not differentiate between higher expression of markers on each EV and a greater percentage of total EVs expressing those markers. CD9, CD63, and CD81 exosome markers were present in all EV groups, although to varying degrees. INF-EV had elevated expression of CD63 (p < 0.0001), while expression of CD9 (p < 0.0001) and CD81 (p < 0.0001) were decreased. Additionally, INF-EV had greatly decreased expression of CD29 (Integrin beta-1, p < 0.0001), CD44 (p < 0.0001), CD49e (Integrin alpha-5, p < 0.0001), CD105 (Endoglin, p < 0.0001), and melanoma-associated chondroitin sulfate proteoglycan (MCSP, p < 0.0001). In contrast, LO2-EV had significantly elevated expression of MCSP (p = 0.0478), CD44 (p = 0.0201), and CD29 (p = 0.0031) compared to NC-EV. LPH-EV did not have significantly different expression of any surface marker compared to NC-EV.

**Figure 2:**
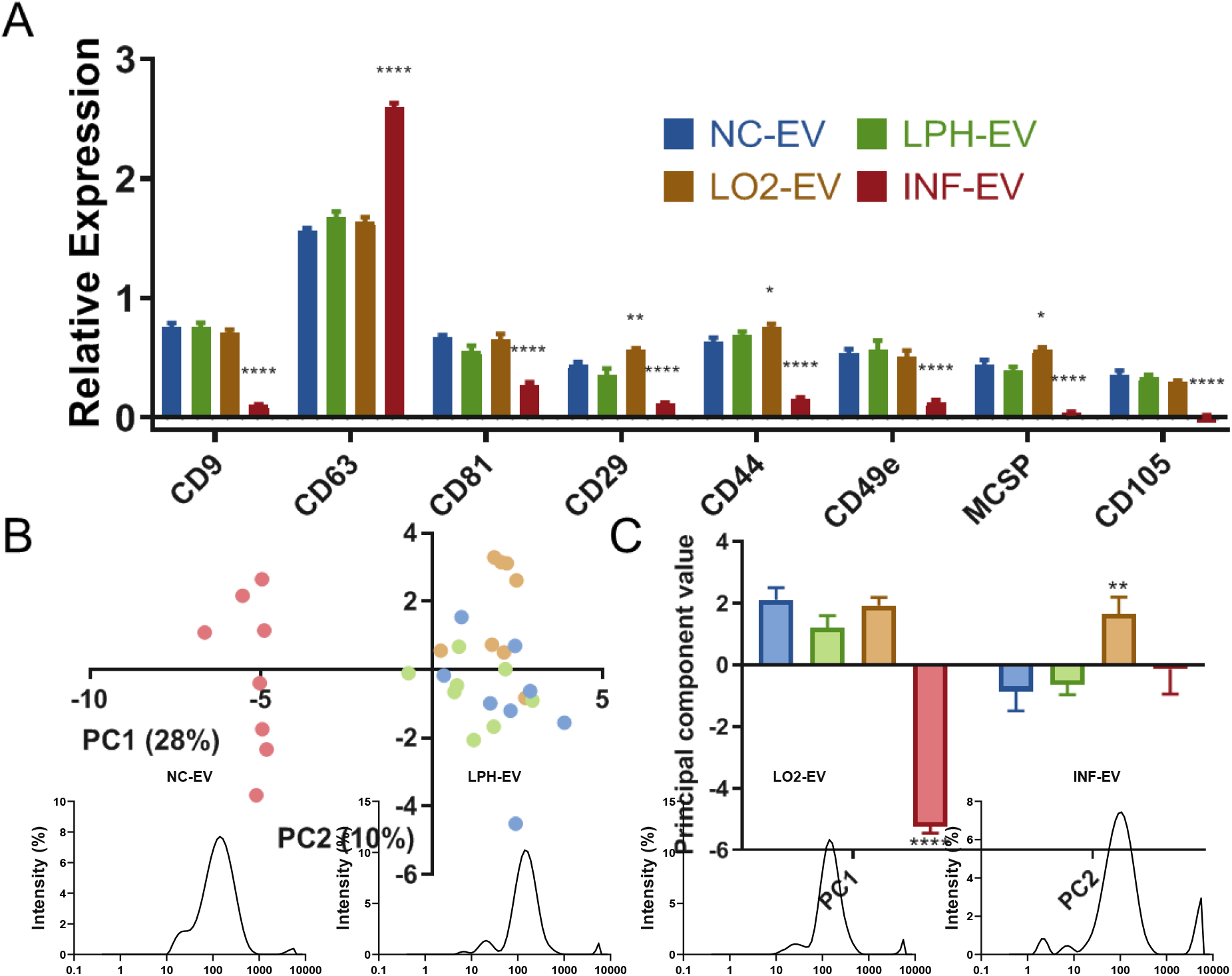
(A) Relative expression of selected EV surface markers across MSC culture conditions as determined by MACSPLEX analysis. MSC priming affects the surface marker expression of released EVs. (B) Plot of PC1 vs PC2 following PCA of all MACSPLEX markers. (C) Plot of PC1 and PC2 values.

Supplementary Figure 4: Dynamic Light Scattering of EVs. Histograms of EV size distribution confirming Two way ANOVA. Data is presented as means ± SEM. N = 3 independent experiments. (*,**, *******) the -existence of a large-diameter population present in INF-MSC-EV at a high concentration. indicate significant difference from Normal Culture at p < 0.05, 0.01, 0.001, 0.0001 by Dunnet’s post hoc test.

PCA enabled visualization of the high dimensional MACSPLEX data to better discriminate differences in the overall expression of surface markers with priming (Fig. 2B). The first and second principal components (PC1, PC2) were responsible for 28% and 10% of the variance in the data set, respectively. INF-EV separated from the other groups along PC1, and INF-EV’s mean value of PC1 was significantly lower than that of NC-EV (p<0.0001, Fig. 2C). Additionally, LO2-EV had a significantly higher mean value of PC2 than NC-EV (p = 0.0018, Fig. 2C). The largest contributors to PC1 were cell adhesion markers, while PC2 was largely made up of cell membrane and immune signaling proteins (Table S5).

**Supplementary Figure 5:**
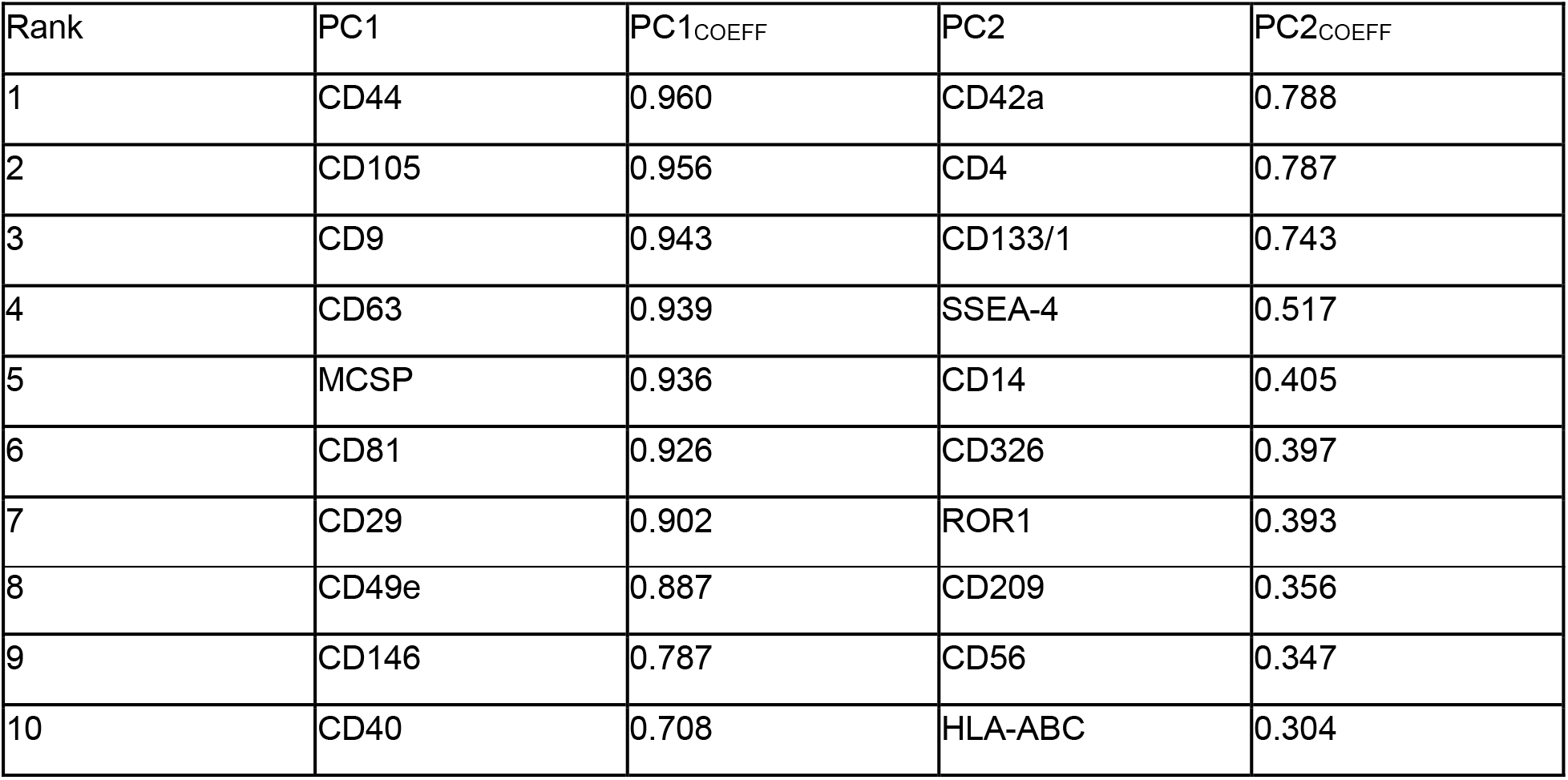
Top 10 contributing surface markers to PC1 and PC2 of MACSPLEX PCA

### 3.3 EV Uptake

CFSE-labeled EV uptake was determined by %CFSE^+^ cells from T cell sub-groups at 24 hours post-treatment. LPH-EV had the highest uptake of all groups (Fig. 3). Treatment with LPH-EV led to significantly greater percent CFSE^+^ cells than the untreated controls for helper T cells (p < 0.05, Fig. 3A), cytotoxic T cells (p < 0.05, Fig. 3B), activated helper T cells, (p < 0.05, Fig. 3C), activated cytotoxic T cells (p <0.01, Fig. 3D), CD4^+^ T_regs_ (Fig. 3E, p < 0.05), and CD8^+^ T_regs_ (Fig. 3F, p < 0.01). No EV group had significant uptake by CD4^−^/CD8^−^ cells (Fig. 3G). LPH-EV was the only EV group to have significant rates of uptake by T cells.

**Figure 3:**
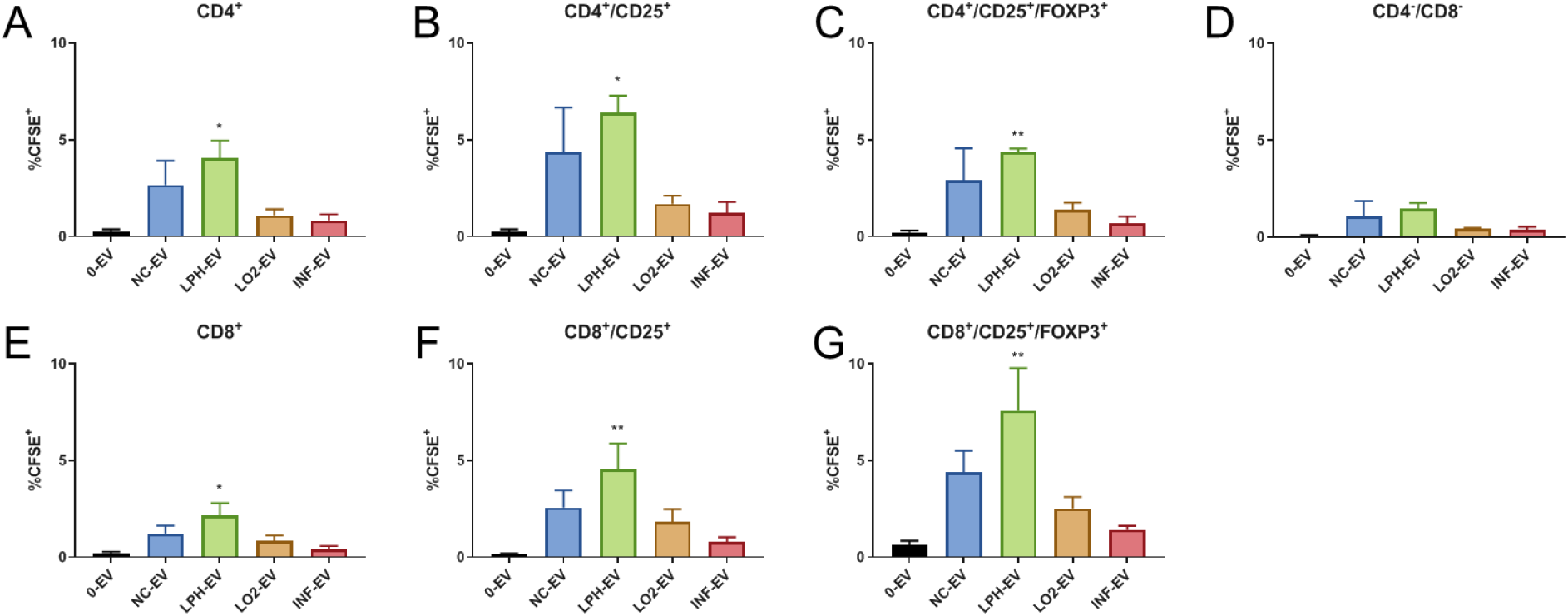
Differential uptake of MSC-EVs by T cell subsets. LPH-EV treated PBMCs had significantly higher CFSE^+^ frequency than untreated cells, indicating uptake of EVs. (A): Frequency of CD4^+^ cells positive for CFSE. (B): Frequency of CD8^+^ cells positive for CFSE. (C): Frequency of CD4^+^/CD25^+^ cells positive for CFSE. (D): Frequency of CD8^+^/CD25^+^ cells positive for CFSE. (E): Frequency of CD4^+^/CD25^+^/FOXP3^+^ cells positive for CFSE. (F): Frequency of CD8^+^/CD25^+^/FOXP3^+^ cells positive for CFSE. (G): Frequency of CD4^−^/CD8^−^cells positive for CFSE. Data is presented as means ± SEM. N = 3 independent experiments. (*,**, ***,****) indicate significant difference from 0-EV at p < 0.05, 0.01,0.001, 0.0001 by Dunnett’s post-hoc test of one-way ANOVA.

### 3.4 Suppression of T cell activation by MSCs and MSC-EVs

PBMCs were subjected to flow analysis to determine T cell activation after 5 days incubation with either MSCs or MSC-EVs. Proliferation of CD8^+^ cells after 5 days was significantly decreased when co-cultured with INF-MSC (p < 0.0001), LO2-MSC (p < 0.0001), LPH-MSC (p < 0.0001), and NC-MSC (p < 0.0001) (Fig. 4B). Similarly, CD4^+^ cell proliferation was significantly decreased by INF-MSC (p = 0.0156), LO2-MSC (p = 0.0191), LPH-MSC (p = 0.0367), and NC-MSC (p = 0.0412) (Fig. 4A). There was no effect on T cell proliferation after 5 days by any MSC-EV group. CD25 expression in helper T cells was significantly reduced by INF-MSC (p = 0.0003), LO2-MSC (p = 0.0050), LPH-MSC (p = 0.0156), and NC-MSC (p = 0.0092) (Fig. 5A). CD25 expression was similarly reduced in cytotoxic T cells cultured with INF-MSC (p = 0.0036), LO2-MSC (p = 0.0238), LPH-MSC (p = 0.0142), and NC-MSC (p = 0.0116) (Fig. 5B). LPH-EV treatment also decreased cytotoxic T cell CD25 expression (p = 0.0218) The frequencies of both CD8^+^ T_regs_ and CD4^+^ T_regs_ were increased by LPH-EV (p = 0.0218, p = 0.0055, respectively). There was no significant change in Treg composition for the MSC only groups (Fig. 5C-D). To determine whether MSC-EVs exhibited time dependent changes in T cell activation, we profiled T cell subset activation following 24 hours of EV treatment and found no significant differences (S5). All MSC groups trended towards a decrease in TNF-α and IFN-γ expression in both helper T cells and cytotoxic T cells, but there were no significant changes (Fig. 5E-H). Overall, MSC co-culture resulted in significant reduction of T cell activation as measured by proliferation and CD25 expression, regardless of priming condition. On the other hand, MSC-EVs had a less pronounced effect with the exception of increased T_reg_ frequency and decreased cytotoxic T cell activation due to LPH-EV treatment.

**Figure 4:**
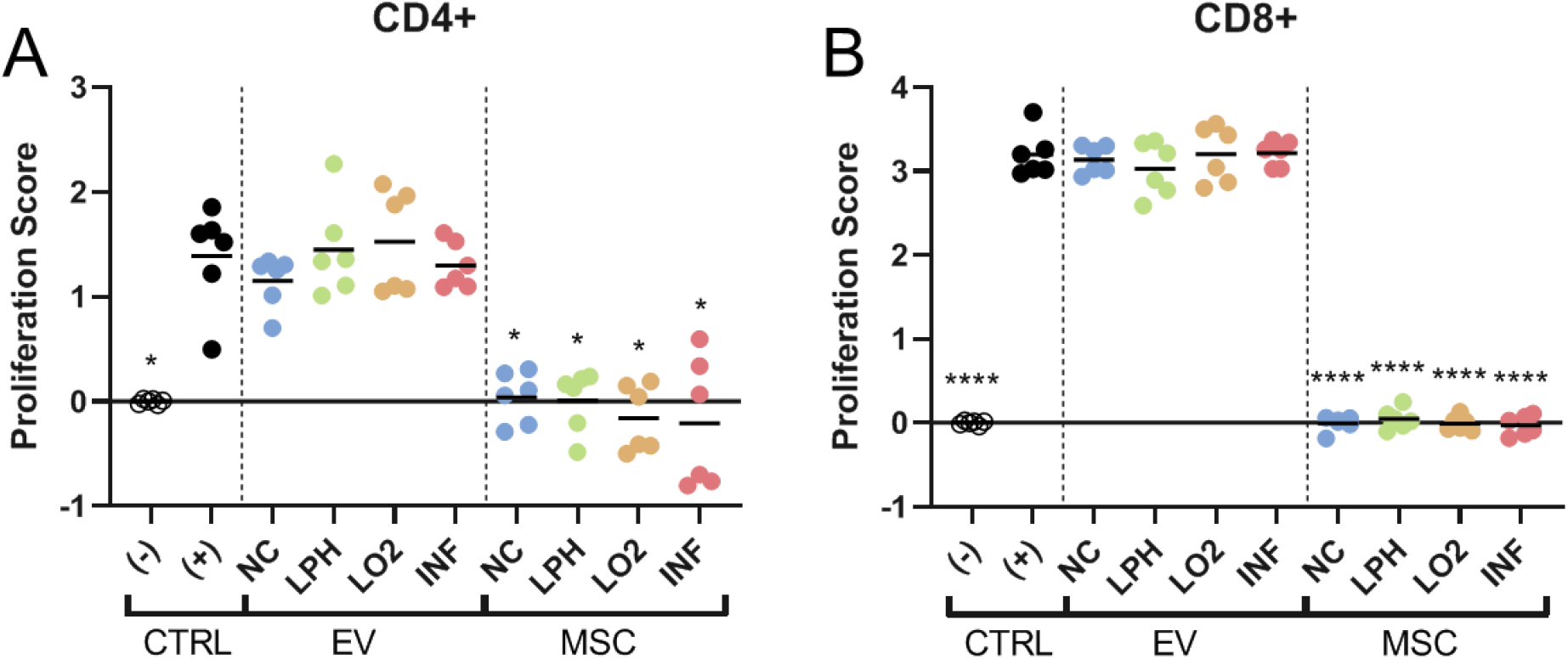
Inhibition of T cell proliferation 5 days after treatment with EVs or MSCs as measured by CFSE dilution. EVs did not affect the proliferation of T cells, while MSCs greatly decreased it. (A): CD4^+^ cells. (B): CD8^+^ cells. Data of two independent experiments consisting of three technical replicates each are presented as means ± SEM. (*,**, ***,****) indicate significant difference from (+) CTRL at p < 0.05, 0.01, 0.001,0.0001 by Dunnett’s post-hoc test of nested one-way ANOVA.

**Figure 5:**
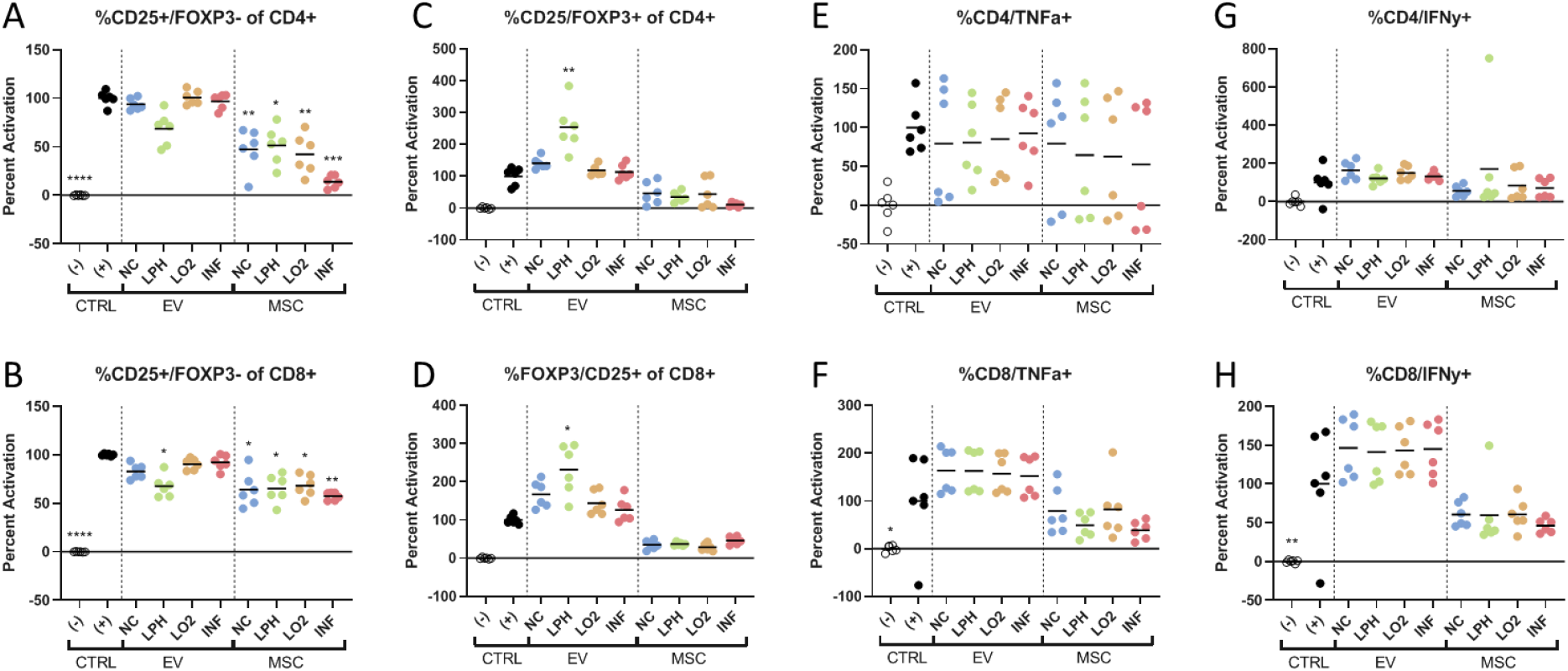
Comparative activation of T-cell subsets 5 days after treatment with EVs or MSCs. Overall, MSCs had significant effects on T effector cells frequency, while LPH-EVs had significant effects on Treg frequency. (A): Frequency of CD4^+^/CD25^+^/FOXP3^−^ cells. (B): Frequency of CD8^+^/CD25^+^/FOXP3^−^ cells. (C): Frequency of CD4^+^/CD25^+^/FOXP3^+^ cells. (D): Frequency of CD8^+^/CD25^+^/FOXP3^+^ cells. (E): Frequency of CD4^+^/TNF-α^+^ cells. (F): Frequency of CD8^+^/ TNF-α^+^ cells. (G): Frequency of CD4^+^/IFN-γ^+^ cells. (H): Frequency of CD8^+^/ IFN-γ ^+^ cells. Data of two independent experiments consisting of three technical replicates each are presented as means ± SEM. (*,**, ***,****) indicate significant difference from (+) CTRL at p < 0.05, 0.01,0.001, 0.0001 by Dunnett’s post-hoc test of nested one-way ANOVA.

**Supplementary Figure 5:**
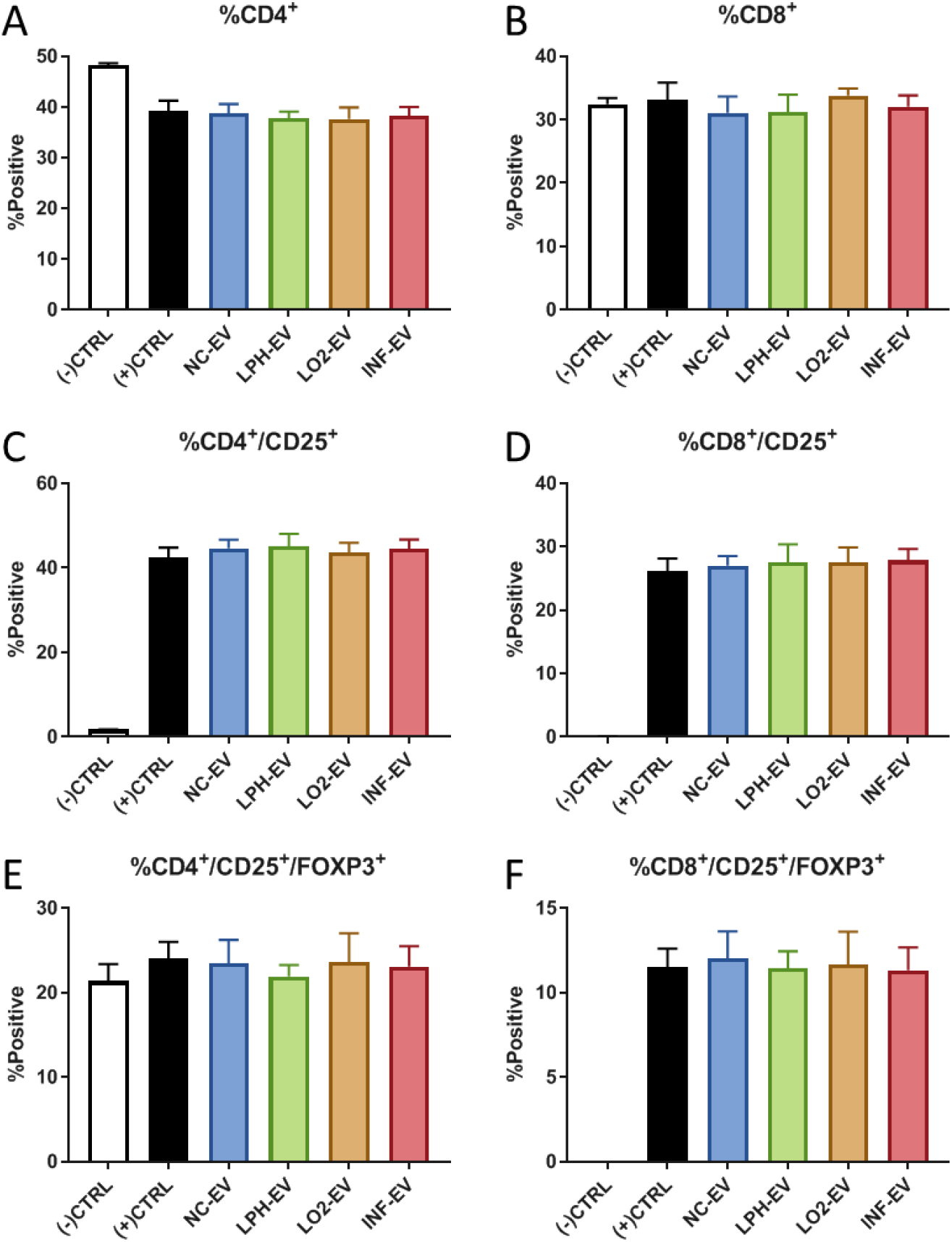
Comparative activation of T cell subsets 24 hours after treatment with CFSE-EVs. There were no significant differences between treatment groups. (A): Percent CD4^+^ cells. (B): Percent CD8^+^ cells. (C): Percent CD25^+^ of CD4^+^ cells. (D): Percent CD25^+^ of CD8^+^ cells. (E): Percent FOXP3^+^ of CD4^+^/CD25^+^ cells. (F): Percent FOXP3^+^ of CD8^+^/CD25^+^ cells. Data is presented as means ± SEM. No significant differences from (+) CTRL group by Dunnett’s post-hoc test of one-way ANOVA.

## 4. Discussion

Although significant evidence exists for the immunomodulatory roles of MSCs and MSC-EVs, the mechanisms of action governing these therapeutic functions are still largely unknown. There is increasing evidence that the MSC microenvironment has substantial effects on the function of their released EVs *in vitro* (21,22). We investigated the effects of different aspects of the injury microenvironment on MSCs and their released EVs. We found that while the yield, size, and surface marker composition of released EVs varied substantially with priming treatments, only those from acidosis primed MSCs had any immunomodulatory effects. These results further emphasize the effect of disease-relevant microenvironment cues on MSCs and could inform the development of future MSC-EV therapeutics.

To our knowledge, this is the first study showing changes in MSC-EV immunomodulation through acidic priming. However, the effect of environmental pH on MSCs has previously been investigated regarding their interactions with various cancers. Tumors are known to create an immunosuppressive microenvironment, and it is hypothesized that MSCs might be involved in this process (34–36). MSCs cultured in an acidic environment enhanced *in vivo* melanoma growth, partly through their increased expression of TGF-β (37). TGF-β is a potent growth factor and has been shown to induce the maturation of T_regs_ (38). MSC-EV have been shown to associate with TGF-β, and EVs may bind TGF-β on their surface (39, 40). Acidosis primed MSCs upregulated osteosarcoma expression of CXCL5 and CCL5 (41). These chemokines have also been implicated in the formation and recruitment of T_regs_, respectively (42–44). MSC-EV may play a role in the MSC driven immunosuppressive role since we observed increased T_regs_ in PBMC cultures treated with acidic preconditioned MSC-EVs.

EV biogenesis and release is known to take place through several mechanisms, including sphingomyelinases (45). Sphingomyelinase activity has been shown to be increased in an acidic environment (46, 47). While increased EV release from MSCs under hypoxic and inflammatory conditions have previously been demonstrated, to our knowledge acidosis has not been previously shown to increase EV release from MSCs (21, 22). Our study describes the first instance of increased EV release by MSCs in an acidic environment, as well as further demonstrating the effect of other priming strategies such as hypoxia and cytokine stimulation.

It was evident that inflammatory priming of MSCs resulted in production of distinct EV populations from other priming conditions. NC-EV, LPH-EV, and LO2-EV had average sizes consistent with exosomes, which range from 50-150 nm. By comparison, INF-EVs exhibited a biomodal size distribution that included a population with size range similar to microvesicles, which bud from the plasma membrane and are between 100nm and 1μm in diameter (45, 48). EVs from both IFN-γ and TNF-α/IFN-γ preconditioned MSCs have previously been observed with larger size distributions tending towards the microvesicle size range (22, 49). The biogenesis of microvesicles differs from that of exosomes, and their immunomodulatory potency has been shown to be less than that of exosomes (50). Additionally, we observed decreases in adhesion-related markers of INF-EV compared to NC-EV. This included integrins beta-1 and alpha-5, as well as endoglin, which is an auxiliary receptor for TGF-β (51). The broad reduction in cell adhesion marker expression of INF-EV was especially evident when viewing the first principal component of the PCA, which was largely composed of these markers (Supplemental Figure 5). Although the MACSPLEX assay lacks a reference standard for absolute quantification, results are comparable within an experiment using equal numbers of EVs (28). The dimensionality reduction of PCA aids in drawing out overall trends in surface marker expression. Cellular recognition and uptake of EVs is regulated in part by their surface markers, which can vary based on the environment of their source cells (22). If INF-EV contain multiple heterogeneous vesicle populations as indicated here, this could dilute the overall potency, possibly explaining the lack of immunomodulation by INF-EVs compared to other studies.

We observed the highest uptake of MSC-EVs in T cells compared to non-T cells across all EV groups (Figure 3). Our results agree with other studies in which MSC-EVs delivered to PBMCs consistently associated with T cells compared to macrophages or NK cells (52). Our study found very little MSC-EV uptake by CD4^−^/CD8^−^ cells. However, MSC-EVs were reported to be primarily taken up by monocytes (22) (49). Previous studies investigated MSC-EV uptake by subsets of immune cells but not the differential uptake by effector and regulatory T cells. The greater uptake of LPH-EV by T_regs_ may be related to their subsequent increased frequency. The decreased expression of cell signaling and adhesion markers in INF-EV may contribute to the observed low uptake for that priming condition.

We observed significantly greater suppression of T cells when they were cultured in direct contact with MSCs versus indirect culture using MSC-EVs. This is in line with previous studies, which have indicated that MSCs interact with T cells differently than their isolated EVs (22, 52). EV dosing may play a role in this, although the dose used in our study, 2000 EV/PBMC, is higher than studies reporting T cell suppression by MSC-EVs (52, 53). As we observed, MSCs are more effective than their EVs alone at inhibiting T cell proliferation (22, 50, 52, 53). However, inhibition of EV release impairs the suppression of T cell proliferation by MSC co-culture, so EVs likely play some role in this process (22). Isolated MSC-EVs also induce T_reg_ formation, while their source cells do not (50, 52), which occurred for the LPH-EV group in this study. Interestingly, there were no significant differences in T cell suppression between any of the direct contact MSC/PBMC co-culture priming conditions. As activated PBMCs create an inflammatory environment of their own, shown here by their production of TNF-α and IFN-γ (Figure 5), it could be that any effects from the initial priming conditions prior to co-culture were overridden by cytokines and other signals produced by the PBMCs.

This study was focused on differences in the release, uptake, and surface composition of EVs; however, we did not look at other aspects of EV potency such as their intravesicular nucleic acid content. MSC-EVs can contain a wide range of micro RNAs (miRNAs), many of which are associated with angiogenesis and tissue remodeling (54). Certain miRNAs have been shown to have increased frequency in EVs derived from MSCs primed with inflammation-relevant signals. When MSCs were primed with TNF-α and IFN-γ, miRNA-155, previously implicated in immune modulation, was increased in their EVs compared to non-primed control MSC-EVs (22, 55). Similarly, IL-1β priming of MSCs upregulated miR-146a, which has previously been shown to regulate the T cell response through the NF-κB pathway (56, 57). MSCs cultured in hypoxia have increased miR-223, miR-146b, miR126, and miR199a (21). Of these, miR-223 is involved in driving anti-inflammatory macrophage polarization, and miR-146b has been shown to monocyte inflammation (58). Further studies will determine whether upregulation of T_regs_ by LPH-EVs is due to change in their miRNA content.

EV biomanufacturing remains an emerging field with enormous potential; however, the lack of standardized methods for characterization and processing further compounds functional heterogeneity observed for MSCs. Differences in manufacturing conditions such as harvesting, isolation, and purification can result in loss of EV subpopulations, exposure of EVs to different stresses, and result in final products with significant heterogeneity (59). For example, EVs frozen at −80°C after isolation and thawed before use have been found to decrease in immunomodulatory potency (50). It will be important to optimize these processing methods to enable proper assessment of EV properties and comparison of EV studies.

## 5. Conclusion

Contributors to EV release and function are still being explored. This study adds to a growing body of evidence demonstrating that EV immunosuppressive function can be enhanced by priming MSCs with inflammation-relevant microenvironment signals. However, there is not yet a consensus on which signals significantly impact EV function, and the mechanisms of action through which MSC-EVs exert their potential therapeutic effects. Comparison of the miRNA, protein, and lipid cargo of EVs from different priming methods could be a future avenue to examine any possible mechanisms for their variable immunosuppressive potential, as well as comparison of MSC-EVs produced by MSCs derived from different donors and tissue sources. Here, MSCs exposed to an acidic environment produce EVs with anti-inflammatory function (i.e. promotion of T_regs_), which could hold great potential to both our understanding of EVs and their eventual clinical translation.

## 6. Acknowledgements

NSF CBET-0939511 and NSF EEC-1648035 funded this work. Julie Nelson and the University of Georgia CTEGD Cytometry Shared Resource Laboratory provided equipment time and expertise. Samantha Spellicy developed the modified EV-CFSE staining protocol used here. Viviana Martinez assisted with DLS analyis of EVs.

